# Preferential recognition and antagonism of SARS-CoV-2 spike glycoprotein binding to 3-*O*-sulfated heparan sulfate

**DOI:** 10.1101/2020.10.08.331751

**Authors:** Vaibhav Tiwari, Ritesh Tandon, Nehru Viji Sankaranarayanan, Jacob C. Beer, Ellen K. Kohlmeir, Michelle Swanson-Mungerson, Umesh R. Desai

## Abstract

The COVID-19 pandemic caused by SARS-CoV-2 is in immediate need of an effective antidote. Although the Spike glycoprotein (SgP) of SARS-CoV-2 has been shown to bind to heparins, the structural features of this interaction, the role of a plausible heparan sulfate proteoglycan (HSPG) receptor, and the antagonism of this pathway through small molecules remain unaddressed. Using an *in vitro* cellular assay, we demonstrate HSPGs modified by the 3-*O*-sulfotransferase isoform-3, but not isoform-5, preferentially increased SgP-mediated cell-to-cell fusion in comparison to control, unmodified, wild-type HSPGs. Computational studies support preferential recognition of the receptor-binding domain of SgP by 3-*O*-sulfated HS sequences. Competition with either fondaparinux, a 3-*O*-sulfated HS-binding oligopeptide, or a synthetic, non-sugar small molecule, blocked SgP-mediated cell-to-cell fusion. Finally, the synthetic, sulfated molecule inhibited fusion of GFP-tagged pseudo SARS-CoV-2 with human 293T cells with sub-micromolar potency. Overall, overexpression of 3-*O*-sulfated HSPGs contribute to fusion of SARS-CoV-2, which could be effectively antagonized by a synthetic, small molecule.

## Introduction

The 2019 novel coronavirus (2019-nCoV, official name: SARS-CoV-2), the causative agent behind the current pandemic, is proving to be highly lethal. SARS-CoV-2 is a member of the family of coronaviruses that generally cause routine infections in humans; however, the severity of organ failure, especially the lung, caused by this virus necessitates studies on all molecular pathways that may be targeted for intervention (1). Of these, virus attachment and internalization pathways are the key to devising strategies that prevent infection even in the early or asymptomatic phase.

The cellular entry of SARS-CoV-2 has been shown to depend on the binding of the viral spike glycoprotein (SgP) to host cell ACE-2 receptor (2-4). Whereas the SgP–ACE-2 pathway has been the focus of most studies, host cell surface heparan sulfate proteoglycans (HSPGs) have also been shown to play important roles in pathology of enveloped viruses, e.g., coronaviruses, herpes simplex virus (HSV), cytomegalovirus, dengue virus, and hepatitis E virus (5-9). In fact, the Esko group has recently shown that infectivity of the SARS-CoV-2 depends on cell surface HSPGs (10). Further, evidence has also been presented that the molecular diversity of HS chains plays an important role in supporting entry, trafficking and replication processes (5), which span a majority of the cellular processes in the life cycle of the virus.

HSPGs contain one or more heparan sulfate (HS) chains covalently linked to serine residues of a core protein such as syndecan and glypican (11). HS is made up of alternating *D*-glucuronic acid (plus some *L*-iduronic acid) and *N*-acetylglucosamine residues that are variably modified by a cascade of sulfotransferases (STs) that engineer sulfated microdomains along the polymeric chain. These microdomains form unique sites of binding for different cell surface receptors, soluble proteins and enzymes (12,13).

A classic ST is the 3-*O*-sulfotransferase, for which six different isoforms (i.e., 3OST-1, -2, -3_A_, -3_B_, -4 and -5) are known (14). Each of these 3OSTs exhibit subtle differences in substrate specificities, thereby engineering rare sulfation microdomains or recognition “codes” in HSPGs (15). While these 3-*O*-sulfate “codes” are significantly different from the common, non-3-*O*-sulfated regions on the HS biopolymer, the selectivity of protein recognition between the multiple 3-*O*-sulfate “codes” may not necessarily be exquisite, as exemplified by the observation that all 3OSTs, except for 3OST-1 (16), support HSV-1 entry and spread (17-21).

Based on the role of HSPGs in viral adherence and internalization (1,21), we hypothesized that SARS-CoV-2 may exhibit subtle role of HSPG microstructure, i.e., the local microdomains within HS, to advantageously gain entry into a host cell. Here, we demonstrate using a model cellular cell-to-cell fusion assay that the SgP of SARS-CoV-2 demonstrates better recognition of 3-OST-3_B_-modified heparan sulfate (HS) receptor in comparison to either the wild-type, unmodified HSPG or the 3OST-5 modified HSPG. Computational studies offer a structural foundation to this role as originating from selective binding of 3-*O*-sulfated sequences to the SgP. More importantly, this pathway could be antagonized by deploying specific agents such as fondaparinux, an anti-3-O-sulfated-HS peptide or a synthetic non-sugar, sulfated, small molecule. Overall, this work provides critical recognition and antagonism insights that should assist developing therapeutics against SARS-CoV-2 attachment and internalization, which could rapidly reduce infection rates.

## Results and Discussion

To assess whether HSPGs mediate cell fusion with an SgP-bearing cell, a luciferase reporter gene activation assay was used in Chinese hamster ovary (CHO-K1) cells, as described in our work (17,22,23). We selected CHO-K1 cells for our cell fusion experiment because these cells lack functional surface receptors for SARS-CoV including human ACE-2 (24). This makes them resistant to infection. Further, CHO-K1 cells are also known to lack endogenous expression of 3-OSTs, which implies that their HSPGs do not carry 3-O-sulfated microdomains (5). In combination, CHO-K1 cells provide an excellent platform to identify new receptor targets that may support SARS-CoV-2 entry in absence of ACE-2.

The cell susceptibility of SARS-CoV-2 being probed here utilizes a cell-to-cell fusion model in which fusion between “effector” and “target” CHO-K1 cells results in activation of the luciferase gene, which is quantified 24h post co-culture using luminescence spectrophotometry. The “effector” cells are transiently co-transfected with plasmids expressing the SARS-CoV-2 SgP and T7 polymerase, while the “target” cells are co-transfected with plasmids carrying either human 3OST-3_B_, 3-OST-5 or ACE-2 genes along with luciferase expression plasmid. As wild-type negative controls, the “target” CHO-K1 cells are co-transfected with an empty vector (pCAGGS), devoid of the 3OST-3_B_, 3-OST-5 or ACE-2 genes, whereas the positive control “target” cells expressed only the human ACE-2 receptor (see Supplementary Materials).

The advantage of this *in vitro* model system is that it affords direct insight into structural features of the SgP–HSPG interaction in cellular settings without the complexities that abound full virus studies. It specifically provides information on receptors other than ACE-2, which may facilitate virus internalization/fusion, especially with regard to modified forms of HS. It is important to note that this cell-to-cell fusion assay is not directed to conclude on the mode of SARS-CoV-2 entry, i.e., membrane fusion or endocytosis, which varies depending on the type of cell (25).

As shown in Figure 1A, the “target” negative control CHO-K1 cells expressing wild-type HS displayed only marginal fusion with “effector” CHO-K1 cells expressing SgP. In contrast, nearly 3-fold higher fusion was detected in the presence of either ACE-2 and/or 3-*O*-sulfated HS receptors. This suggests a critical role for the 3OST-3_B_ modified HSPG receptor in SgP-mediated cell-to-cell fusion during viral spread. It is important to note that 3OST-3_B_-modified HSPG “target” cells do not contain the well-established SARS-CoV-2 receptor ACE-2 (26-28). Further, transfecting varying levels of 3OST-3_B_ plasmid alone into the effector cells led to a concomitant increase in cell-to-cell fusion (see Figure S1). Thus, the results show that SgP-mediated cell-to-cell fusion arises even in the absence of ACE-2.

**Figure 1.**
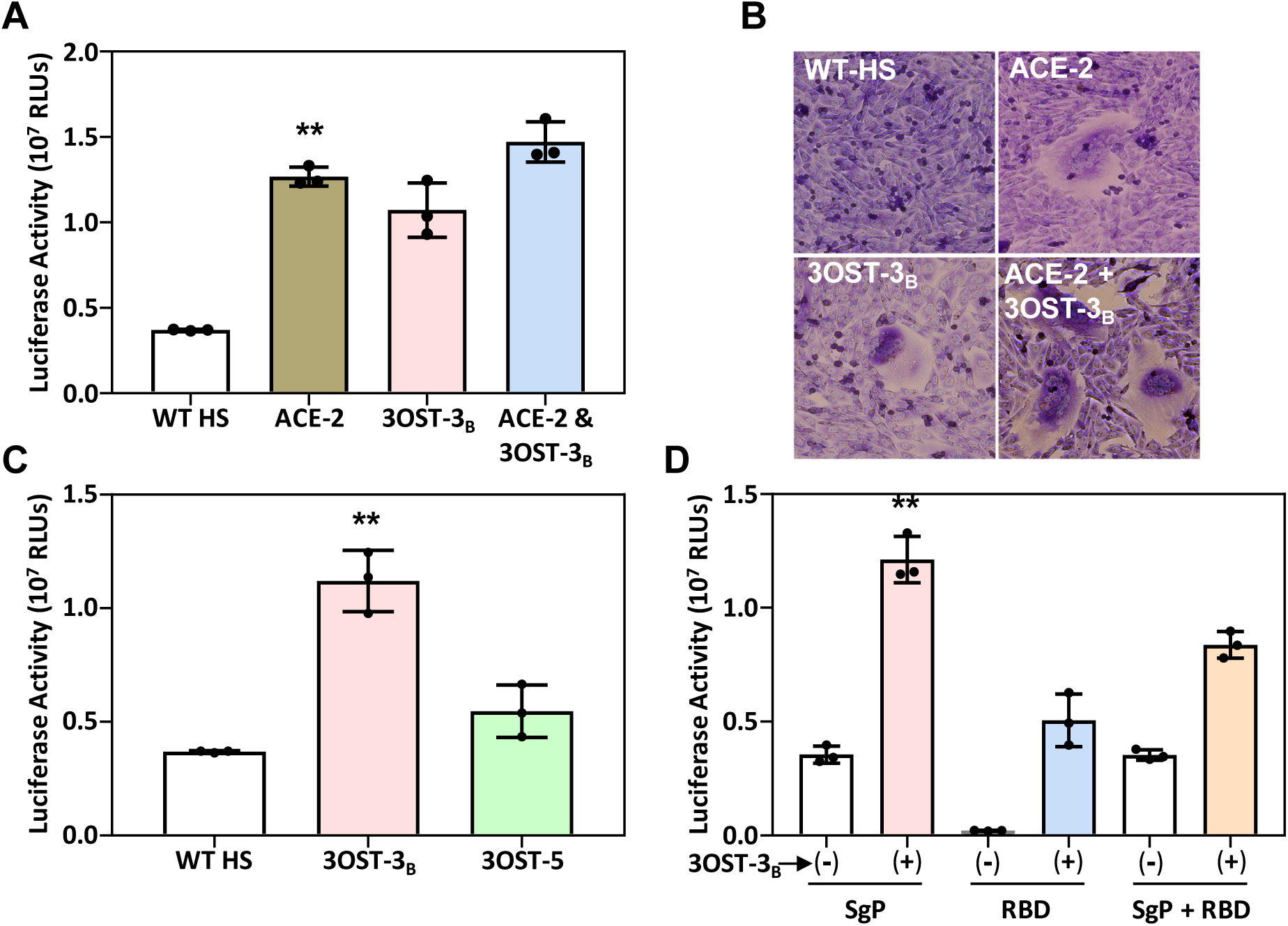
Selective recognition of 3-O-sulfated heparan sulfate (HS) by SgP of SARS-CoV-2 in promotion of cell fusion. (A) “Effector” CHO-K1 cells expressed SgP of SARS-CoV-2, while the “target” CHO-K1 cells expressed 3OST-3_B_ or ACE-2. Wild-type (WT) HS refer to “target” negative control CHO-K1 cells expressing neither 3OST-3_B_ nor ACE-2 receptor. (B) Microscopic visualization of syncytia induced by SgP–HSPG or SgP–ACE-2 interaction. (C) Comparison of cell fusion when “target” HSPGs carry either 3OST-3_B_ or 3OST-5 generated sequences. Error bars = ±1 SD; ** P<0.05, one-way ANOVA.

Interestingly, no statistical difference in cell-to-cell fusion between CHO-K1 cells carrying either ACE-2 or 3OST-3_B_ HSPG receptors was observed (Figure 1A). Additionally, the combined presence of ACE-2 and 3-*O*-sulfated HSPG did not exhibit any additivity or synergism under the conditions studied. Although this suggests saturation of cell-to-cell fusion with either 3-*O*-sulfated HSPGs or ACE-2-mediated pathways, detailed studies would be needed to ascertain absence of synergism under all conditions.

The SgP of SARS-CoV-2, through specific interactions with the host cell receptor, is also known to contribute to syncytia formation (29), an important pathway in the pathogenesis of the virus. Therefore, we next assessed syncytia formation by co-culturing SgP expressing “effector” cell with “target” CHO-K1 cells expressing HSPGs or ACE-2. As shown in the Figure 1B, minimal (or no) syncytia formation was observed for target cells carrying wild-type HSPG receptor (Panel ‘WT-HS’), while much higher number of syncytia were observed in SgP-mediated fusion with target cells carrying either ACE-2 (Panel ‘ACE-2’) or HSPGs modified by 3OST-3_B_. Here, co-expression of both 3OST-3_B_ and ACE-2 displayed higher syncytia numbers. Taken together, these results ascertain that 3-*O*-sulfated HSPGs are likely to play an important role in SgP-mediated cell-to-cell fusion process.

We next assessed selectivity of 3OST-modified sub-domain recognition by SARS-CoV-2 SgP by comparing target CHO-K1 cells transfected with plasmids expressing either 3OST-3_B_ or 3OST-5. The results showed that cells expressing 3OST-3_B_ displayed ∼3-fold more cell-to-cell fusion than 3OST-5 expressing cells (Figure 1C). In fact, fusion with 3OST-5 effector cells was very similar to the negative control, wild-type HSPG-carrying effector cells. This implies that SARS-CoV-2 SgP differentially recognizes the 3-*O*-sulfated HS structures generated by the two different isoforms, 3OST-3_B_ and 3OST-5. This result is strikingly different from HSV-1, which can utilize both 3OST-3 and 3OST-5 isoform-generated HSPGs for cell fusion (17). Thus, it appears that SgP of SARS-CoV-2 is more selective in its recognition of the HSPG receptor. A quick question to address here is whether differential expression of 3OST-3 and 3OST-5 could have contributed to differential recognition of SgP. To mitigate this possibility, we performed simultaneous experiments for HSV-1 entry. Our results suggested that cells expressing both 3OST-3_B_ and 3OST-5, but not the control cells (i.e., containing unmodified HSPGs), allowed HSV-1 entry (5,17) (see Figure S2). Finally, we also tested whether more relevant human cells, i.e., human lung epithelial A549 cells, would exhibit 3-OST-3_B_ enhanced SgP–mediated cell fusion. Figure S3 shows that 3OST-3_B_ overexpression enhances fusion in the manner observed for CHO-K1 cells. Thus, SARS-CoV-2 SgP recognition of HSPGs is different from that HSV-1 glyroproteins.

To better understand SgP features that contribute to recognition of HSPGs and mediation of cell-to-cell fusion, we studied full length SgP as well as its receptor-binding domain (RBD) alone. Full length SgP contains two subunits, S1 and S2. The RBD is located within the S1 subunit (2,31,32) and is known to mediate interaction with the ACE-2 receptor. In contrast, the S2 subunit contributes to cellular internalization (33,34). Do these functions hold when 3-*O*-sulfated HSPG is the only receptor present on target cells?

To address this, the effector cells expressing either full length SgP, the RBD alone or both were co-cultured with target cells expressing the 3OST-3_B_ modified HSPG or negative control target cells expressing wild-type HSPG. The results indicated that full length SgP, but not RBD alone, promoted 3-*O*-sulfated HSPG pathway (Figure 1D). Alternatively, in the absence of the ACE-2 receptor on target cells, fusion with effector cells occurs only with full-length SgP and is dramatically impaired with RBD alone. This implies that the S2 subunit is critical for cell-to-cell fusion mediated by the SgP–3-*O*-sulfated HSPG pathway. More importantly, cell fusion was miniscule when the effector cells expressing RBD alone were exposed to wild-type HSPG target cells in comparison to moderate level of cell fusion for 3-*O*-sulfated HSPG target cells (Figure 1D). A similar phenotype with no cell fusion was observed when ACE-2 target cells were co-cultured with effector cells carrying RBD alone (see Figure S4). This implies that the RBD preferentially recognizes 3-*O*-sulfated microdomains generated by 3OST-3_B_.

We next addressed the question whether this selectivity of SgP recognition originates from differences in atomistic interactions with different sequences of HS. Recent studies have shown that the selectivity of HS recognition can be predicted through rigorous a dual-filter computational algorithm that compare and contrast interactions of a large number of HS sequences binding to proteins (35). We built a library of 27,930 unique topologies possible for natural di-, tetra, and hexasaccharides of HS in a combinatorial manner. This library of HS sequences was studied for recognition of the RBD of SgP trimer. Each of the 27,930 topologies was docked in triplicate using a genetic algorithm-based using a well-established dual-filter algorithm (Figure 2A). All three plausible sites of HS binding on SgP trimer, proposed recently by Linhardt and coworkers (36), were studied (Figures 2B and S5). Of these, the RBD was found to be the most favored site for HSPG to interact (not shown).

**Figure 2.**
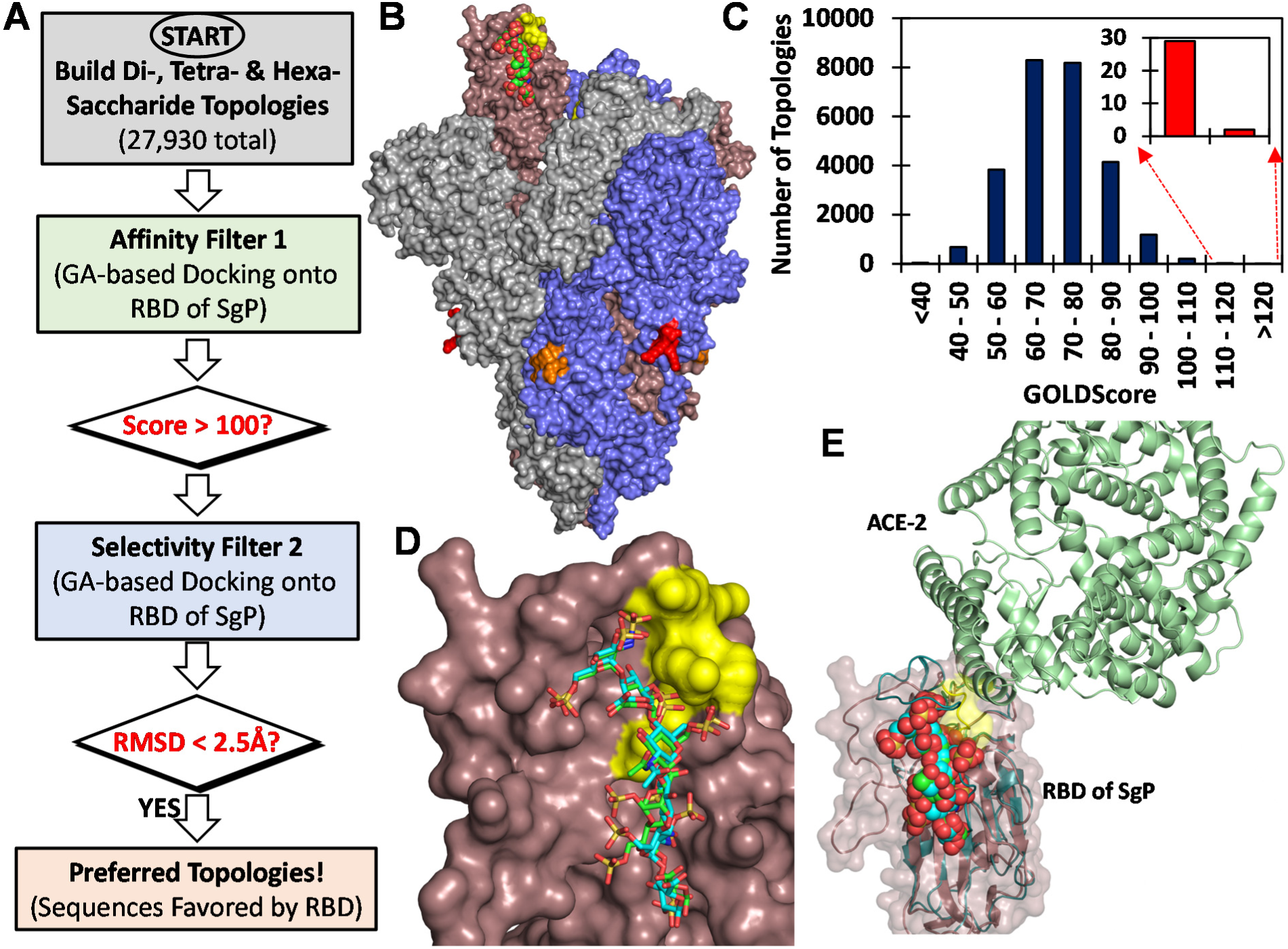
Computational screening of a library of di-, tetra- and hexa-saccharide sequences of HS (total 27,930 topologies) against RBD of SgP for identification of origin of selectivity at the atomistic level. (A) The plausible sites of HS binding onto SgP trimer include ^453^YRLFRKS^459^ (yellow = RBD), ^681^PRRARS^686^ (red); and ^810^SKPSKRS^816^ (orange) in the trimeric SgP (chains A, B, and C are shown in pink, grey and blue, respectively). (B) The dual-filter algorithm used to identify high affinity and high specificity HS topologies that bind to the RBD of SgP. GOLDscore was the first filter, while RMSD (consistency of binding) was the second filter. (C) Results after the first filter in the form of a histogram of the number of HS hexasaccharide topologies for every 10 unit change in GOLDscore. Inset shows promising high affinity topologies. (D) The zoomed version of the two high selectivity HS hexasaccharides binding to the RBD of SgP (sticks in green and cyan color). See Figure S4 for details on the structure of these sequences and their nature of interaction with residues constituting the RBD of SgP. (E) Overlay showing the two HS hexasaccharides (shown in van der Waals rendering) binding in the RBD domain of SgP (trimer) in relationship with the known ACE2 (green ribbons) site of binding from the cryo-EM structure (PDB:6M0J).

The predicted poses of HS binding onto RBD were analyzed using two parameters corresponding to *in silico* affinity (GOLD score) and consistency of binding (i.e, root mean square deviation (RMSD)) as implemented in our studies on multiple protein–HS systems (16,37,38). The analysis indicated that none of the 30 di-or 900 tetra-sequences recognized SgP well (not shown). In contrast, 242 of the 27,000 hexasaccharide topologies were predicted to interact with the RBD with high affinity (Score >100, Figure 2C). Nearly 92% of these, or 223 unique topologies, contained at least one 3-*O*-sulfated glucosamine residue. An unusual structural characteristic of this group was the preferred placement of the 3-*O*-sulfate group on either the 1^st^ or 3^rd^ residue from the non-reducing end (see Table S1). In fact, majority of HS sequences favored by SgP contained multiple 3-*O*-sulfated residues suggesting a structural characteristic that is likely to be unique for SgP.

Interestingly, two sequences displayed high affinity as well as high selectivity for RBD (Score > 120, RMSD < 2.5 Å; Figure 2D) by forming strong ionic and hydrogen bonding interactions (see Figure S6). Residues common in both sequences included GlcNS3S6S, IdoA2S and GlcNAc6S, which engineered high selectivity of interaction (Figure 2E). Interestingly, a glucuronic acid is present on the non-reducing end of the 3-*O*-sulfated glucosamine in one sequence, which is not known to be generated by 3OST-5 (39). Thus, these computational studies afford strong atomistic foundation to the concept that the RBD of SgP preferentially recognizes 3-*O*-sulfated microdomains of HS.

We reasoned that the selectivity of SgP for 3-*O*-sulfated HS could offer a route to discovering antagonists of cell-to-cell fusion. To study this, we first tested whether treatment with bacterial heparinase I, which can partially degrade cell surface HSPGs, would decrease HS-dependent fusion of “target” CHO-K1 cells with SgP-bearing “effector” cells. It is important to note that heparinase I prefers heparin as a substrate (40); however, HS is also known to be depolymerized. As shown in Figure 3A, heparinase I-treated target cells displayed ∼2-fold cell-to-cell fusion. Next we considered competitive antagonism, which has a better potential in terms of therapeutics. Hence, we studied fusion between 3-*O*-sulfated HS-bearing “target” cells and SgP-containing “effector” cells in the presence of a generic peptide targeting common HS sequences (G1) and a specific peptide directed towards 3-*O*-sulfated HS (G2). The G1 and G2 peptides were developed earlier, using phage display library screening, as probes for studying HS selectivity against HSV entry and spread (17,21-23). As evident from Figure 3A, 10 µM dose of both peptides depressed cell fusion, as would be expected on the basis of competitive antagonism. However, a striking difference of nearly 2-fold was observed for cells bearing the 3-*O*-sulfated HS receptor in comparison to the wild-type HS receptor (Figure 3A).

**Figure 3.**
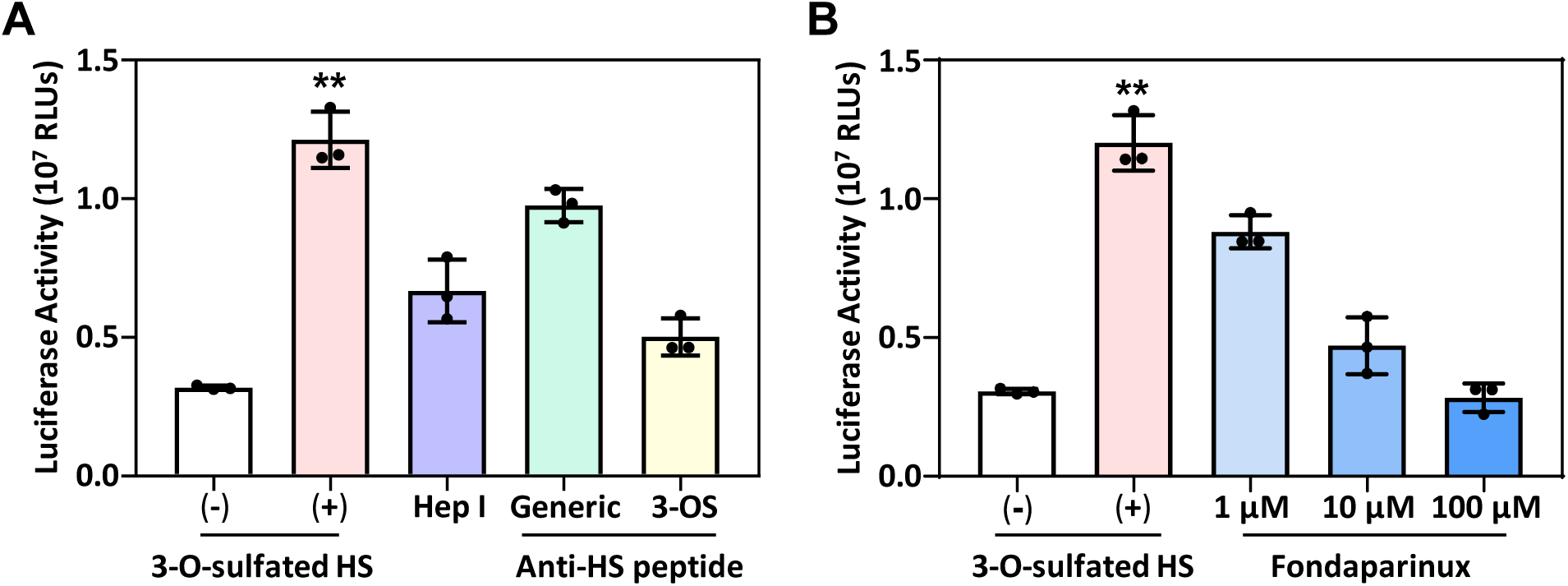
Antagonism of 3-O-sulfated HS receptor through (A) either cleavage of HSPGs by heparinase I (Hep 1, 1.5 units/ml), competitive inhibition with generic (10 µM) or 3-O-sulfate-specific (10 µM) peptides, or (B) competition with 3-O-sulfate containing synthetic pentasaccharide, fondaparinux. “Effector” CHO-K1 cells expressed SgP of SARS-CoV-2, while “target” CHO-K1 cells expressed 3OST-3_B_. Wild-type HS indicated by (-) refer to “target” negative control CHO-K1 cells devoid of 3OST-3_B_ expression. Error bars = ±1 SD; ** P<0.05, one-way ANOVA.

To further test the selectivity of 3-*O*-sulfated HS recognition and antagonism, we utilized fondaparinux, a clinically used anticoagulant. Fondaparinux is a unique, synthetic pentasaccharide with a central 3-*O*-sulfated glucosamine residue. As shown in Figure 3B, the addition of fondaparinux inhibited SgP-mediated cell-to-cell fusion with “target” cells expressing 3-*O*-sulfated HS in a dose-dependent manner. At nearly 100 µM level, fondaparinux reduced cell fusion to the basal levels observed for the wild-type HS receptor. This result further confirms the role of 3-*O*-sulfated HS in mediating SgP-based cell fusion.

The highly sulfated nature of the HS hexasaccharides identified through computational studies (Figures 2 and S6, Table S1), also led us to reason that a synthetic, small, non-sugar, highly sulfated compound, called SPGG, could serve as an effective inhibitor of cell-to-cell fusion mediated by SgP (Figure 4). SPGG has recently been identified as a highly promising pan-virus antagonist of cellular entry because it competes for viral glycoproteins, such as glycoprotein D of HSV, that are involved in recognition of cell surface HSPGs (41-44). We studied SPGG’s effect on cell-to-cell fusion in CHO-K1 cells as well human HEK293T cells. As shown in Figures 4A and 4B, the synthetic agent SPGG reduced cell-to-cell fusion in both cell lines quite effectively. As the concentration of SPGG increased to 1.0 µM, cell-to-cell fusion for both cell lines decreased approximately 50%. This suggested a strong possibility that SPGG may exhibit highly promising anti-SARS-CoV-2 potential.

**Figure 4.**
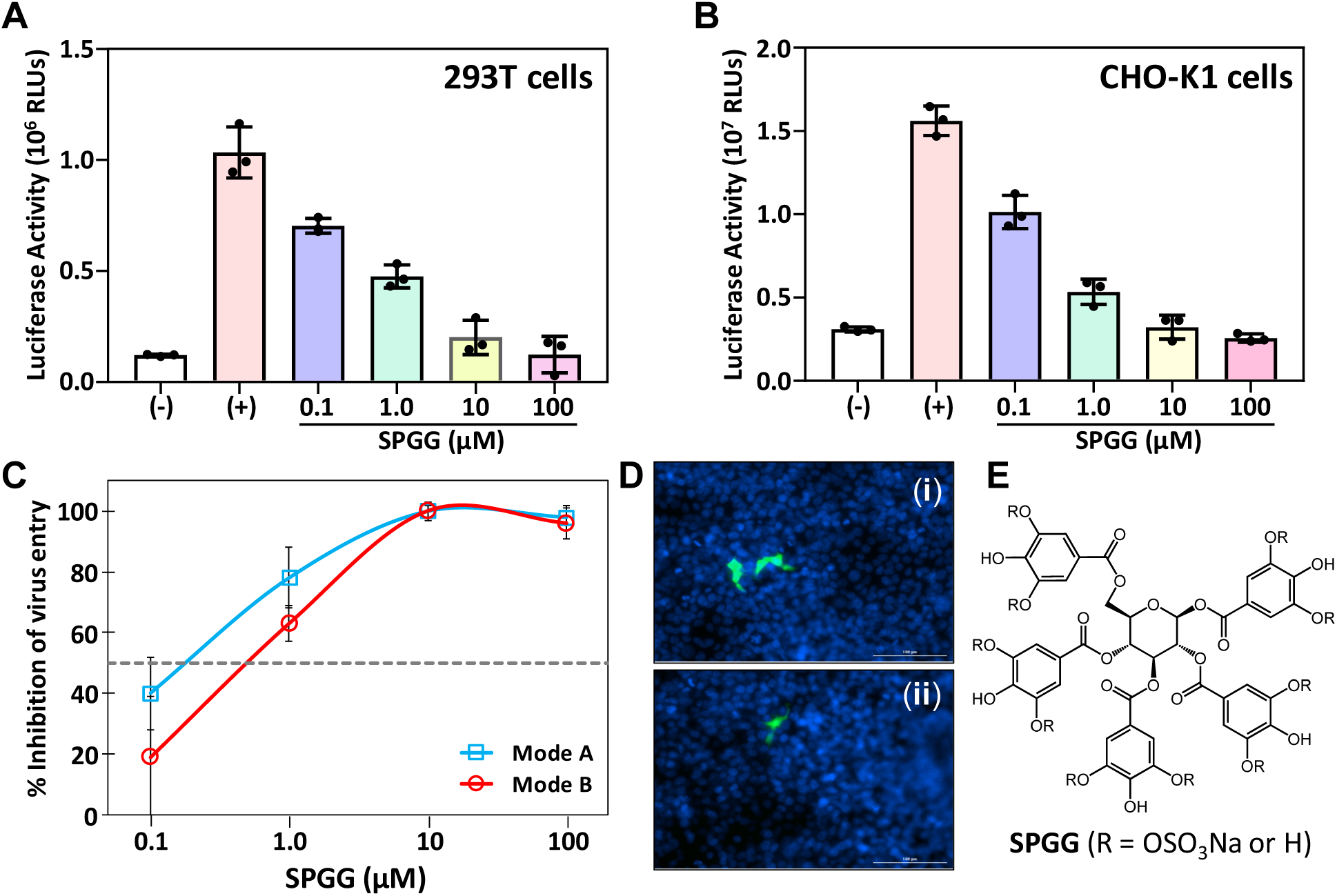
Antagonism of SgP-mediated cell fusion by SPGG, a synthetic, small, highly sulfated compound in two different systems. (A) Inhibition of cell-to-cell fusion between “effector” CHO-K1 cells expressing SgP of SARS-CoV-2 and “target” 293T cells (A) or CHO-K1 cells (B). Wild-type HS expressing in in both cell types represents the negative control (shown as (-)). Positive control 293T cells (shown as (+)) express ACE-2 receptor, while that in the CHO-K1 cells express 3OST-3_B_ enzyme, which modifies surface HSPGs to 3-O-sulfated HS. Error bars = ±1 SD; ** P<0.05, one-way ANOVA. (C) Inhibition of SARS-CoV-2 pseudo typed lentivirus with SPGG. Treatment of either the pseudo-virus (mode A, blue) or 293T cells (mode B, red) with SPGG reduced viral entry (measured 48h post-transduction) using GFP-fluorescence of the pseudo-virus. Error bars = ±1 SD. Grey dotted line represents 50% inhibition. D) Representative fluorescence microscopy of the mock-treated (panel (i)) or SPGG-treated 293T cells (panel (ii)) at 10X magnification. (E) Structure of SPGG.

To further assess the translatability of SPGG’s anti-cell-to-cell fusion activity, we generated GFP-expressing pseudo-typed SARS-CoV-2 particles (pLV-S) using a third-generation lentivirus-system (45). These particles express the SARS-CoV-2 SgP on their surface in the native form. Briefly, HEK293T cells were co-transfected with a *pLV-eGFP* (the GFP plasmid), *psPAX2* (packaging plasmid) and either *pCAGGS-S* (SARS-CoV-2 plasmid) or VSV-G (control plasmid) (see Supplementary Materials). Following 48 h of co-culture, the virus (or control) particles were harvested, quantified and used for infection of new HEK293T cells at dilutions of 10^2^ to 10^7^ so that virus titers were in the range of 20 to 100 GFP positive cells. Two treatment modalities were explored. In mode A), SPGG was pre-incubated with the pLV-S pseudovirus before infecting the target 293T cells. In mode B), the 293T cells were first pre-treated with SPGG and then challenged with the pLV-S pseudo-virus.

Figure 4C shows that SPGG severely impaired viral entry in a dose-dependent manner in both treatment modalities. Interestingly, the modality of treatment did not impact SPGG’s inhibition potential. Most importantly, the inhibition potency of SPGG could be between 0.1 and 1.0 µM (see 50% inhibition line, Figure 4C). A comparison of GFP fluorescence of 293T cells revealed significant decrease in internalized GFP-tagged virus particles in the presence of SPGG (panel (ii), Figure 4D). Overall, the synthetic agent SPGG was identified to be a promising inhibitor of pseudo-typed SARS-CoV-2 entry into human 293T cells.

## Conclusions

In this study, we have provided the first evidence that 3-*O*-sulfated microdomains in HS chains of HSPGs, especially those generated by 3OST-3_B_, but not necessarily by 3OST-5, offer preferential recognition of SgP of SARS-CoV-2. This recognition affords unique opportunities for development of antagonists that may prevent cell-to-cell fusion and viral spread. Of the several antagonists studied in this work, the synthetic, highly sulfated small molecule SPGG was especially effective in reducing SgP-mediated cell-to-cell fusion and syncytia formation. Considering that this pathway has been reported for human enteroids (46) and also serves as a mechanism to evade antibody neutralization (47), SPGG as a sub-micromolar antagonist of this pathway has major translational value.

The entry of SARS-CoV-2 into host cells is a multi-step process involving several molecular and cellular factors. Whereas optimal and timely functioning of each of these factors is needed for the virus to multiply and spread, SgP is an obligatory factor because of its key role in cell surface receptor engagement, without which no virus entry is possible (2). ACE-2 was identified as the first host cell surface receptor for SARS-CoV-2 entry (3,31,32). Yet, growing evidence points to HSPG as another receptor that the virus uses to anchor to the host cell (10). This work establishes that modified forms of HS, especially those produced by 3-OST-3_B_, greatly enhance cell fusion to facilitate host cell entry. This result is of major significance because it has been known that inflamed cells and tissues of the lung exhibit higher levels of 3-*O*-sulfated HS (48). This implies that the inflammatory storm induced by SARS-CoV-2, especially in the lung, is likely to be significantly aided by 3-*O*-sulfated HSPGs, thereby orchestrating a self-feeding, self-destructive cycle with a detrimental outcome.

Our model studies with CHO-K1 “target” and “effector” cells show that 3-*O*-sulfated HS alone can promote cell-to-cell fusion (Figure 1A). Our cell-fusion assay is a focused attempt to identify receptors other than ACE-2, which was clearly evident from the results with the 3-O sulfated modified forms of HS. One advantage of this assay is that functional domains and amino acid residues in SgP can be identified relatively easily to understand the impact of mutational changes. Such studies may be very worthwhile since SARS-CoV-2 has shown variability in the SgPs (2,49).

This suggests that as far as cell-to-cell fusion process (50) is concerned 3-*O*-sulfated HSPGs on host cells may act as functional receptors, independent of ACE-2. Alternatively, other forms of cellular entry, e.g., through fusogenic proteins, may require ACE-2. As evident in recent studies (10), HSPGs play supporting role in such processes.

This work provides a unique insight in SgP recognition of 3-*O*-sulfated HS and ACE-2 receptors. Our work shows that 3OST-3_B_ modified HSPG, but not wild-type HSPG, mediates fusion with the RBD alone (Figure 1D). Interestingly, a recent study has shown that the heparin binding to the RBD is accompanied by conformational changes, which may play a role in cell entry (51). In striking contrast, cell fusion with RBD alone was not observed for the ACE-2 receptor (Figure S4). Rather, it is well established that fusion mediated by the ACE-2 receptor requires cleavage of the full length SgP into S1 and S2 units (2,3). This highlights the differentiating role of 3-*O*-sulfated microdomains of HS in SgP mediated cell-to-cell fusion, especially through the RBD. Further, this work highlights the importance of studying other 3OSTs to elucidate additional receptor requirements, if any.

It is important to note that the activity of HSPG modifying enzymes, e.g., 3OSTs, is dependent on the availability of optimal substrate sites within existing HS chains (15,16,52). In fact, the activity of the cascade of sulfotransferases and their isoforms (NDST, 2OST and 6OST), would determine the generation of 3-*O*-sulfated microdomains. This implies that populations of patients carrying upregulated HS biosynthetic genes may be more susceptible to SARS-CoV-2 infection. Rigorous studies with samples from different types of patients would be needed to evaluate this deduction from our work.

Several studies have shown that heparin or heparin-like molecules bind SgP with high affinity. Studies on full length and low molecular weight heparins by the Linhardt group measured potencies in the picomolar range (36), whereas the Boons group have reported nanomolar potencies for both polymeric and oligomeric heparins (53). Such high potencies could be advantageously translated into potential drug candidates, as proposed by the Turnbull group in the form of a clinical-stage heparan sulfate mimetic (54). Suramin, a small molecule mimetic of heparin, has also been presented as an inhibitor of early steps of SARS-CoV-2 infection (55).

Our work greatly expands these possibilities to include 3-*O*-sulfate microdomain targeting peptides as well as synthetic, high sulfated SPGG. Likewise, fondaparinux, which has a 3-*O*-sulfate group, is also a significant inhibitory agent. Of these, SPGG is particularly attractive because of its ease of synthesis, lack of cytotoxicity, and broad spectrum antimicrobial activity (42,43,56). SPGG is also an inhibitor of coagulation factor XIa (56), which would simultaneously induce an anti-coagulant effect without major bleeding risk. Thus, use of SPGG in SAR-CoV-2 has the potential of addressing episodes of thrombosis observed in large number of severely ill patients (57).

Overall, our results provide the first evidence of a role of 3-O-sulfated HS in SARS-CoV-2 SgP mediated cell fusion and viral entry. HS clearly plays a role in SARS-CoV-2 pathogenesis; however, 3-*O*-sulfated HS, especially when overexpressed, greatly enhances cell fusion. This work lays the foundation for development of small molecule agents against SARS-CoV-2. A good number of HS mimetics are currently in clinical trials, especially against cancer. This work highlights the promise of such HS mimetics in treatment and prevention of SARS-CoV-2 too.

## Acknowledgements

We thank the computational resources made available to us through a grant from the National Center for Research Resources (S10 RR027411) to VCU.

